# Decreased Extracellular Vesicle Vasorin in Severe Preeclampsia Plasma Mediates Endothelial Dysfunction

**DOI:** 10.1101/2024.06.24.600441

**Authors:** Saravanakumar Murugesan, Dylan R Addis, Hanna Hussey, Mark F Powell, Lakshmi Saravanakumar, Adam B Sturdivant, Rachel G Sinkey, Michelle D Tubinis, Zachary R Massey, James A Mobley, Alan N Tita, Tamas Jilling, Dan E Berkowitz

## Abstract

**Background:** Preeclampsia (PE) is a serious pregnancy complication affecting 5-8% of pregnancies globally. It is a leading cause of maternal and neonatal morbidity and mortality. Despite its prevalence, the underlying mechanisms of PE remain unclear. This study aimed to determine the potential role of vasorin (VASN) in PE pathogenesis by investigating its levels in extracellular vesicles (EV) and its effects on vascular function.

**Methods & Results:** We conducted unbiased proteomics on urine-derived EV from severe PE (sPE) and normotensive pregnant women (NTP), identifying differential protein abundances. Out of one hundred and twenty proteins with ≥ ±1.5-fold regulation at P<0.05 between sPE and NTP, we focused on Vasorin (VASN), which is downregulated in sPE in urinary EV, in plasma EV and in the placenta and is a known regulator of vascular function. We generated EV with high VASN content from both human and murine placenta explants (Plex EV), which recapitulated disease-state-dependent effects on vascular function observed when treating murine aorta rings (MAR) or human aortic endothelial cells (HAEC) with murine or human plasma-derived EV. In normal murine pregnancy, VASN increases with gestational age (GA), and VASN is decreased in plasma EV, in placenta tissue and in Plex EV after intravenous administration of adenovirus encoding short FMS-like tyrosine kinase 1 (sFLT-1), a murine model of PE (murine-PE). VASN is decreased in plasma EV, in placenta tissue and in EV isolated from conditioned media collected from placenta explants (Plex EV) in patients with sPE as compared to NTP. Human sPE and murine-PE plasma EV and Plex EV impair migration, tube formation, and induces apoptosis in human aortic endothelial cells (HAEC) and inhibit acetylcholine-induced vasorelaxation in murine vascular rings (MAR). VASN over-expression counteracts the effects of sPE EV treatment in HAEC and MAR. RNA sequencing revealed that over-expression or knock down of VASN in HAEC results in contrasting effects on transcript levels of hundreds of genes associated with vasculogenesis, endothelial cell proliferation, migration and apoptosis.

**Conclusions:** The data suggest that VASN, delivered to the endothelium via EV, regulates vascular function and that the loss of EV VASN may be one of the mechanistic drivers of PE.

**CLINICAL PERSPECTIVE:** What is New

- VASN in circulating plasma EV in sPE is reduced compared with VASN content in plasma EV of gestational age-matched pregnant women.
- VASN is encapsulated and transported in EV and plays a pro-angiogenic role during pregnancy.
- VASN should be explored both for its pro-angiogenic mechanistic role and as a novel biomarker and potential predictive diagnostic marker for the onset and severity of PE.

What Are the Clinical Implications?

- VASN plays a role in maintaining vascular health and the normal adaptive cardiovascular response in pregnancy. A decrease of VASN is observed in sPE patients contributing to cardiovascular maladaptation.
- Strategies to boost diminished VASN levels and/or to pharmacologically manipulate mechanisms downstream of VASN may be explored for potential therapeutic benefit in PE.
- The decrease in EV-associated VASN could potentially be used as a (predictive) biomarker for PE.

## Introduction

Hypertensive disorders during pregnancy, affecting 5-8 % of pregnancies globally, pose lifelong risks, including cardiovascular disease, hypertension, renal failure, stroke, and reduced life expectancy^1^. Preeclampsia (PE) also leads to fetal growth restriction and is a primary cause of medical labor induction and preterm birth, posing significant risks to both mothers and infants^2–4^. Although the exact mechanisms behind PE remain unclear, they involve an interplay among placenta-derived factors, immune responses, and maternal vascular systems, leading to vascular imbalance characterized by vasoconstriction, endothelial leakage, increased inflammation, and poor placentation^5,6^. These mechanisms disrupt the normal proangiogenic and vasodilatory environments, resulting in vascular remodeling and abnormal vascular reactivity. Various signaling pathways are implicated, including VEGF, PlGF, TGF-β, the renin angiotensin system (RAS), autoantibodies against angiotensin and endothelin receptors, complement activation, systemic inflammation, and HIF1α activation ^7–19^.

Extracellular vesicles (EV), membrane-bound vesicles found in biological fluids, are increasingly recognized as critical mediators of intercellular communication in normal pregnancy ^20^. However, placental EV may also contribute to pathological states. Treatment of vascular rings or cultured endothelial cells with sPE EV results in a reduction in endothelial nitric oxide synthase (eNOS) and endothelial dysfunction, a hallmark of PE ^21^. Moreover, studies demonstrate that placental EV isolated from PE patients can induce hypertension in animal models ^22^. While EV are implicated in PE-associated vascular dysfunction, the precise mechanisms remain under investigation. ^21,23–27^. Our current study focuses on the protein cargo of EV, as the role of microRNAs within PE-derived EV has been extensively explored ^26,28–36^.

VASN, a glycosylated type I transmembrane protein structurally similar to the Drosophila melanogaster slit protein, exhibits significant cross-species conservation ^37^. ADAM17 metalloprotease cleavage releases the soluble form of VASN, which acts as a potent inhibitor of TGF-β signaling and modulates angiotensin II (Ang II) signaling ^38^. Reduced VASN can promote arterial aging by insufficient inhibition of ang II/TGF-β fibrogenic signaling^39^. VASN^−/−^ mice develop myocardial mitochondrial injury and cardiac hypertrophy^40^. Additionally, Taggi et al. reported the first evidence of VASN expression in human female reproductive tissues ^41^. Importantly, a previous study demonstrated that VASN downregulation in vascular smooth muscle cells after acute vascular injury contributes to the fibroproliferative response. ^37^ The antagonism of fibroproliferative response by VASN is mediated by the direct binding of VASN’s extracellular domain to TGF-β family members, thereby antagonizing their signaling^38^. This discovery underscores the critical regulatory role of VASN in vascular health and disease. However, further investigation is necessary to elucidate the specific role of VASN in PE and related pathologies.

Previously, we demonstrated that EV cargo isolated from sPE plasma induces endothelial dysfunction and impairs vasorelaxation ^21^. In the current study, we hypothesized that quantitative changes in the protein cargo of sPE EV cause PE pathologies. Utilizing unbiased proteomic analysis of urinary EV, we identified VASN as a promising candidate for mediating vascular changes in PE. Here, we detail our findings as we explore the hypothesis that altered VASN content within circulating sPE EV contributes to the observed impaired vascular endothelial function in sPE.

## Methods

### Data Access

The data that support the findings of this study are available from the corresponding author upon request.

### Patient Population

This study was reviewed and approved by the University of Alabama at Birmingham (UAB) institutional review board (IRB) (IRB-300004235) and written informed consent was obtained from all subjects participating in the study. The study was registered prior to patient enrollment on October 24, 2019, and registered with ClinicalTrials.gov (NCT04154332, Mark Powell, November 6, 2019). Patient recruitment occurred on the labor and delivery unit. All patients admitted to the labor and delivery unit for delivery were eligible for screening. This included patients admitted for induction of labor, those in active labor, and those scheduled for cesarean section. Two groups were recruited for this study: (1) sPE and (2) normotensive. Inclusion criteria for the sPE group were age ≥ 18 years and a diagnosis of PE with severe features^21^ (a) blood pressure ≥ 160/110 mmHg after 20 weeks gestation and ≥ 300 mg/day proteinuria or a protein/creatinine ration of ≥ 0.3 mg/mg; or (b) blood pressure ≥ 160/110 mmHg after 20 weeks gestation with any of the following co-conditions: platelet count less than 100,000 X 10^9^/L, AST/ALT enzymes elevated to twice the upper limit of normal, serum creatinine >1.1 mg/dL or a doubling of the creatinine from baseline, pulmonary edema, new-onset headache, and/or visual disturbances. Inclusion criteria for the normotensive group were age ≥ 18 years and no diagnosis of chronic or pregnancy-associated hypertension. After consent was obtained, samples (blood, urine, and placenta) were collected from the patients. The urine samples were collected with protease inhibitors, centrifuged at 1600 x g for 10 minutes, and stored at −80°C for further analysis. Prior to delivery, 10 mL of whole blood was collected via venipuncture in an anticoagulant ethylenediaminetetraacetic acid–K2 (EDTA-K2) tube and centrifuged at 3000 x g for 20 minutes. With respect to the timing of the whole blood collection, the only requirements were for the blood to be collected after the patient was committed to delivery and before the delivery of the newborn. Blood was routinely drawn with scheduled lab work during labor for patient convenience. For cesarean deliveries, the blood was drawn after placement of the neuraxial block.

### Murine model of PE

Experiments using mice were approved by the Institutional Animal Care and Use Committee of the University of Alabama at Birmingham (IACUC-21860). Adenoviral constructs (Ad-CMV-hsFlt-1 and Ad-CMV-GFP) are injected into the retroorbital sinus of pregnant mice at a dose of 2.5×10^9^ PFU in 100μL volume on E11.5, and then the injections are repeated on E15.5. *<u>Monitoring, measurements and samples:</u>* 1) Measurements of body weights daily. 2) Terminal procedures at E18.5: a) Collection of blood under anesthesia. b) Collection of aorta for wire myography, and aorta, liver, and placenta for RNA and protein isolation and histological examination. c) Assessment of fetal weights and fetal survival.

### Isolation of EV from urine samples

At time of collection, 10mL of urine was aliquoted into a 15-mL conical tube with appropriate amount of protease inhibitors (e.g., 100uL of HALT protease inhibitor cocktail, 100x, PI78429, Fisher Scientific) and centrifuged at 1000×g for 10min at 4°C. The supernatant was then be stored at −80°C until further processing. A day before the EV enrichment, urine was thawed overnight at 4°C. On the day of enrichment, samples were centrifuged first at 17K×g for 10min at 4°C using a fixed angle rotor. Supernatant was collected in a fresh 50-mL conical tube (Supernatant1, or SN1). The pellet was then be resuspended in 0.5mL of 0.5M Urea, pH 7.6, with 200mg/mL final concentration of DTT and incubated at 37°C for 10min (with vortexing every 2 min). Dulbecco’s phosphate-buffered saline (DPBS) was added to the final volume of 10mL, and samples was centrifuged at 17k×g for 10min at 4°C in the fixed angle rotor. The supernatant (SN2) was combined with SN1 in the 50-mL conical tube. The SN1 SN2 mixture were centrifuged in a swing bucket rotor at 200K×g for 1hr at 4°C. The supernatant was collected in a separate 50-mL conical tube and stored at −80°C in case any analysis of EV-free urine is needed in future. The 200×g centrifugation process in the same ultracentrifuge tube was repeated until all SN1 SN2 mixture is pelleted. The pellet was be washed with 8-10mL of DPBS 1-2 time(s) via centrifugation at 200K×g for 1hr at 4°C. Discard the washes. The pellet was resuspended in 30uL of 1xLDS sample buffer (NP0007, Fisher Scientific), sonicated in an ultrasonic water bath for 20min, quantified using an EZQ protein assay. Samples was reduced with NuPAGE Reducing Reagent (NP0009, Fisher Scientific), denatured at 70°C for 10min, and separate onto a 10% Bis-tris gel (NP0315BOX, Fisher Scientific, run short-stack: ∼1cm from the top). The gel was stained using a Colloidal Blue Staining kit (LC6025, Fisher Scientific) following the manufacturer’s instruction. Samples were further processed for MS/proteomics analysis at this point.

### Placenta collection

This study utilized human term placentas (37–39 weeks gestation) obtained from both normotensive pregnant women (NTP) and women diagnosed with severe preeclampsia (sPE) with approval from the University of Alabama at Birmingham, Human Research Ethics Committee (IRB-300004235). Written informed consent was obtained from all participants prior to placenta collection, which occurred immediately following normal labor or elective cesarean delivery. The collected placentas were subsequently used to establish placental explant cultures (Plex) for further experimentation.

### Placental explant culture

Following delivery, term placental tissue samples (15-18 g) were obtained from both NTP (n = 6) and sPE groups. Villous explants were dissected from the decidual layer at a depth of 2-3 mm. These explants were thoroughly washed with sterile phosphate-buffered saline (PBS) to remove residual blood. Subsequently, the villous tissues were meticulously diced into small pieces (∼2 mm³) and distributed into 100 mm culture dishes (approximately 25 pieces per dish) containing 30 mL of serum-free RPMI 1640 medium (Roswell Park Memorial Institute medium). To achieve further dissociation, the tissue pieces were gently pressed through a sterile 200-micron stainless-steel mesh using a glass pestle. The resulting cell suspension was then layered onto a Percoll gradient for red blood cell (RBC) depletion. The enriched cell fraction, collected from the top of the Percoll layer, was washed and plated in fresh RPMI medium. These cultures were then incubated for 24 hours at 37°C in a humidified incubator with 95% air and 5% CO_2_. Following incubation, the conditioned media was collected for subsequent EV isolation using precipitation methods. ^21^

### Isolation of EV from placental explant cultures

EV were isolated from the conditioned media of placental explant cultures (Plex) using the ExoQuick-TC isolation kit (System Bioscience, Inc.) according to the manufacturer’s instructions^22^ Briefly, 5 ml of Plex media was thoroughly mixed with 1 ml of ExoQuick-TC solution (ExS) and incubated overnight at 4°C. The mixture was then centrifuged at 1500×g for 15 minutes at 4°C to pellet the EVs. The resulting EV pellets were resuspended in Dulbecco’s phosphate-buffered saline (DPBS) and stored at −80°C until further analysis (10). following minimal information for studies of Extracellular Vesicles (MISEV) 2023 guidelines ^42^.

### Proteomics analysis

Proteomics analysis was followed by previous publications ^43–46^. In brief, the protein fractions were quantified using an 8-point Pierce BCA Protein Assay Kit (Thermo Fisher Scientific, PI23225), and 20 µg of protein per sample diluted to 35µL using NuPAGE LDS sample buffer (1x final conc., Invitrogen, NP0007). Proteins were then reduced with Dichlorodiphenyltrichloroethane (DTT) and denatured at 70°C for 10 min prior to loading onto Novex NuPAGE 10% Bis-Tris Protein gels (Invitrogen, NP0315BOX) and separated half way (15min at 200 constant voltage). The gels were stained overnight with Novex Colloidal Blue Staining kit (Invitrogen, LC6025). Following de-staining, each lane was partitioned into three separate MW fractions and equilibrated in 100mM ammonium bicarbonate (Millipore SIGMA; 1066-33-7). Each gel plug was then digested overnight with Trypsin Gold, Mass Spectrometry Grade (Promega, V5280) following manufacturer’s instruction. Peptide extracts were reconstituted in 0.1% Formic Acid/ and double distilled H_2_O at 0.1µg/µL.

Peptide digests (8µL each) were injected onto a 1260 Infinity nHPLC stack (Agilent Technologies), and separated using a 100-micron I.D. x 13.5 cm pulled tip C-18 column (Jupiter C-18 300 Å, 5 microns, Phenomenex). This system runs in-line with a Thermo Orbitrap Velos Pro hybrid mass spectrometer, equipped with a nano-electrospray source (Thermo Fisher Scientific), and all data were collected in collision induced dissociation (CID) mode. The nHPLC was configured with binary mobile phases that included solvent A (0.1%FA in ddH2O), and solvent B (0.1%FA in 15% ddH2O / 85% ACN), programmed as follows; 10min @ 5%B (2µL/ min, load), 90min @ 5%-40%B (linear: 0.5nL/ min, analyze), 5min @ 70%B (2µL/ min, wash), 10min @ 0%B (2µL/ min, equilibrate). Following each parent ion scan (300-1200m/z @ 60k resolution), fragmentation data (MS2) was collected on the top most intense 15 ions. For data dependent scans, charge state screening and dynamic exclusion were enabled with a repeat count of 2, repeat duration of 30s, and exclusion duration of 90s.

The XCalibur RAW files were collected in profile mode, centroided and converted to MzXML using ReAdW v. 3.5.1. The data was searched using SEQUEST, which was set for two maximum missed cleavages, a precursor mass window of 20ppm, trypsin digestion, variable modification C @ 57.0293, and M @ 15.9949. Searches were performed separately with both Human and Rat species-specific subsets of the UniProtKB databases. Following this analysis, since our goal was to compare the human proteome data for this study, we utilized the high-confidence rattus norvegicus to human tryptic peptide homolog data to avoid confusion for down-stream Systems Analysis.

The list of peptide IDs generated based on SEQUEST (Thermo Fisher Scientific) search results were filtered using Scaffold (Protein Sciences, Portland Oregon). Scaffold filters and groups all peptides to generate and retain only high confidence IDs while also generating normalized spectral counts (N-SC’s) across all samples for the purpose of relative quantification. The filter cut-off values were set with minimum peptide length of >5 aminoacids (AA), with no (*m/z*+1) charge states, with peptide probabilities of >80% confidence intervals (C.I.), and with the number of peptides per protein ≥2. The protein probabilities were then set to a >99.0% C.I., and a Dlaw False Discovery Rate (FDR)<1.0. Scaffold incorporates the two most common methods for statistical validation of large proteome datasets, the false discovery rate (FDR) and protein probability (28; 42; 56). Relative quantification across experiments were then performed via spectral counting(35).

For the proteomic data generated, two separate non-parametric statistical analyses were performed between each pair-wise comparison. These non-parametric analyses include 1) the calculation of weight values by significance analysis of microarray (SAM; cut off >|0.6|combined with 2) t-test (single tail, unequal variance, cut off of p < 0.05), which then were sorted according to the highest statistical relevance in each comparison. For SAM (15) whereby the weight value (W) is a statistically derived function that approaches significance as the distance between the means (μ1-μ2) for each group increases, and the SD (δ1-δ2) decreases using the formula, W=(μ1-μ2)/(δ1-δ2). For protein abundance ratios determined with N-SC’s, we set a 1.5-2.0-fold change as the threshold for significance, determined empirically by analyzing the inner-quartile data from the control experiment indicated above using ln-ln plots, where the Pierson’s correlation coefficient (R) was 0.98, and >99% of the normalized intensities fell between +/−1.5 fold. In each case, any two of the three tests (SAM, t-test, or fold change) had to pass.

### Western Blot Analysis

EV suspended in DPBS were prepared by addition of radio immunoprecipitation assay buffer. Total protein concentration was measured using the bicinchoninic acid assay (Pierce; Thermo Fisher Scientific, Bonn, Germany). Proteins were separated and transferred to polyvinylidene fluoride membranes by gel electrophoresis and electro blotting, respectively. The blots were incubated with EV specific antibodies (VASN, TSG101, AGT and β-Actin (1:5000; CST, CA, USA) antibodies overnight at 4°C. Then the blots were incubated for 1 hour at room temperature with a goat anti-rabbit– horseradish peroxidase antibody (1:5000; CST). Membranes were incubated with the chemiluminescent substrate and visualized using the ChemiDoc®Gel Documentation System (Bio-Rad Laboratories, Hercules, CA, USA). ^21^

### RNA isolation and analysis

25 mg tissue sections were bead-lysed in RLT buffer (Qiagen) supplemented with 1% (w/w) 2-mercaptoethanol and frozen. RNA was isolated from cells with RNeasy kit with Qiashredder cell disruption (Qiagen) and RNase-free DNase Set (Qiagen) on-column DNA digestion. RNA purity and concentration were assessed using an Agilent 2100 Bioanalyzer RNA Analysis chip (Eukaryote Total RNA Pico Series II) (Agilent Technologies). Quantitative real-time PCR (qPCR) was performed using a Rotor-Gene Q instrument (Qiagen), TaqMan MGB primer/probe sets (Thermo Fisher) and Premix Ex Taq master mixer (Takara).

### Cell Migration Assay

To assess the effect of EV from NTP, sPE women and an EV-free control on HAEC, cells were cultured in 6-well plates. Wounds were made by scraping a conventional pipette tip across the cell monolayer. The wound images were captured by microscopy (AmScope) immediately after wounding and after 24 hours. The multiple wound width was measured using the ImageJ software, and the percentage of wound healed was calculated using the following formula: Wounded area filled (%) = 100% − (width after 24 hours/width at the beginning) × 100%.

### Transmission Electron Microscopy (TEM) and Nanoparticle Tracking Analysis (NTA)

For morphological characterization, Plex EV were adsorbed onto continuous carbon grids and negatively stained with 2% uranyl acetate solution. The size and morphology of the isolated EV were subsequently visualized using transmission electron microscopy (TEM) at an accelerating voltage of 120 kV at the UAB High-Resolution Imaging Facility (HRIF). Additionally, the size distribution of the Plex EV was determined using nanoparticle tracking analysis (NTA) employing a NanoSight NS300 instrument (HRIF, UAB) analysis. ^21^

### Immunofluorescence labeling of EV uptake in endothelial cells

EV dual labeling assays were performed as we described in previous publications. ^21^ In brief, Triton X-100 (0.001 %) was used to permeabilize the EV. Then, ExoQuick precipitation solution (EPS-100µl) was added to precipitate the EV pellets and centrifuged at 3000 x g at room temperature (RT) for 5 minutes. The supernatant was carefully removed, and the pellets were dissolved in DPBS (100μl). 1μg of TSG101 (ab83; Abcam, USA), AGT and VASN (ab156868: Abcam, USA) antibody were added to the EV suspension in a microfuge tube and incubated for 1.5 hours with gentle shaking at room temperature (RT). EPS (100µl) was added and centrifuged at 3000 x g for 5 minutes. Again, the pellets were resuspended in 100µl of DPBS. The above EPS step was repeated twice to remove the excess antibody. The precipitated EV pellets were resuspended in 100µl of DPBS. Then, Alexa Fluor labeled secondary (Anti-mouse IgG, 4409, Cell Signaling Technology, USA) antibody (1:100 dilution) was added to the EV suspension and incubated for 60 minutes with gentle shaking in the dark. EPS (100µl) was added to EV suspension and centrifuged at 3000 x g for 5 minutes, 3 times. The pellets were dissolved in 50 µl of PKH diluent and add 0.2µl of PKH67 (PKH67GL, Sigma, USA). Next, the EV suspension mixture was incubated for 30 minutes with gentle shaking. The EPS (50µl) was added to EV suspension mixture and centrifuged at 3000 x g for 5 minutes. The EV pellets were dissolved in 50µl of DPBS and 300μl EV-depleted serum was added to stop the reaction. The EPS (100µl) was added to EVs suspension mixture and centrifuged at 3000 x g for 10 minutes. The EV pellets were then dissolved with 20µl of EV-depleted medium and incubated with HAEC for 24 hours. After 24hours, the cells were washed with DPBS, fixed with 4% paraformaldehyde for 30 minutes at RT, and were stained in blue with DAPI (1; 5000; Invitrogen; Thermo Fisher Scientific, Inc), PKH67 (green-excitation at 488 nm) and EV (red-excitation at 594 nm) for 5 minutes. After washing with DPBS, the cells were photographed using a confocal microscope (Nikon TE2000 inverted microscope, HRIF, UAB) and analyzed with Nis Elements software.

## Statistical Analysis

Results were analyzed with 1- or, when appropriate, 2-way ANOVA for multiple comparisons using GraphPad Prism, version 8.01, followed by Tukey’s post hoc test to adjust for multiple comparisons. Data are presented as the mean ± SEM, with all data points shown as scatter. All experiments were performed on at least six biological replicates (n= 6); the specific number of replicates are reported for each experiment. Two group means were analyzed by Student’s t-test. Significance levels were determined between samples examined and were set at p<0.05. Statistical analyses were performed using Prism 8 (GraphPad Inc, CA, USA).

## RESULTS

### Unbiased proteomics performed on EV isolated from urine of NTP and sPE patients

Previously, we reported that plasma EV from sPE mediated endothelial dysfunction ex vivo^21^. To identify potential protein cargo molecule candidates mediating this effect, we performed unbiased proteomic analysis to identify differentially expressed proteins between urinary exosomes isolated from NTP and women with sPE. Urinary EV were chosen for this discovery phase of the project due to the relative ease to obtain plasma protein contamination-free EV from urine, as opposed to plasma. Demographic data of the patient population used for this analysis can be found in **Supplemental Table 1.** Principal component analysis (PCA) demonstrated distinct clustering of NTP and sPE samples along PC1, explaining 50% of the variance **(Figure 1A).** A heatmap with hierarchical clustering further confirmed the separation of the NTP and sPE groups based on their overall protein contents **(Figure 1B).** A total of 321 unique proteins were identified in EVs, with 121 exhibiting significant differential regulation (fold-change > 1.5, p-value < 0.05) between the two groups. We ranked significantly regulated genes based on their Cohen’s d standardized effect size from largest to smallest **(Figure 1C)**. **Figure 1D** shows a heatmap of the top 20 highest regulated (in absolute terms; up or down) ranked from top to bottom. We highlighted VASN as one of the highest regulated proteins between sPE and NTP, which is downregulated in sPE and its role in pregnancy or in PE is unknown. Disease ontology (DO) analysis using overrepresentation analysis (ORA) and gene set enrichment analysis (GSEA) revealed significant enrichment of pathways related to vascular, kidney, and hepatic diseases, including PE, among the differentially regulated proteins **(Figure 1D).** A gene concept network was constructed to visualize the relationships between these proteins and the enriched DO pathways **(Figure 1E).** The color of each dot in the network represents the relative change in protein abundance (blue for decrease, magenta for increase).

**Figure 1.**
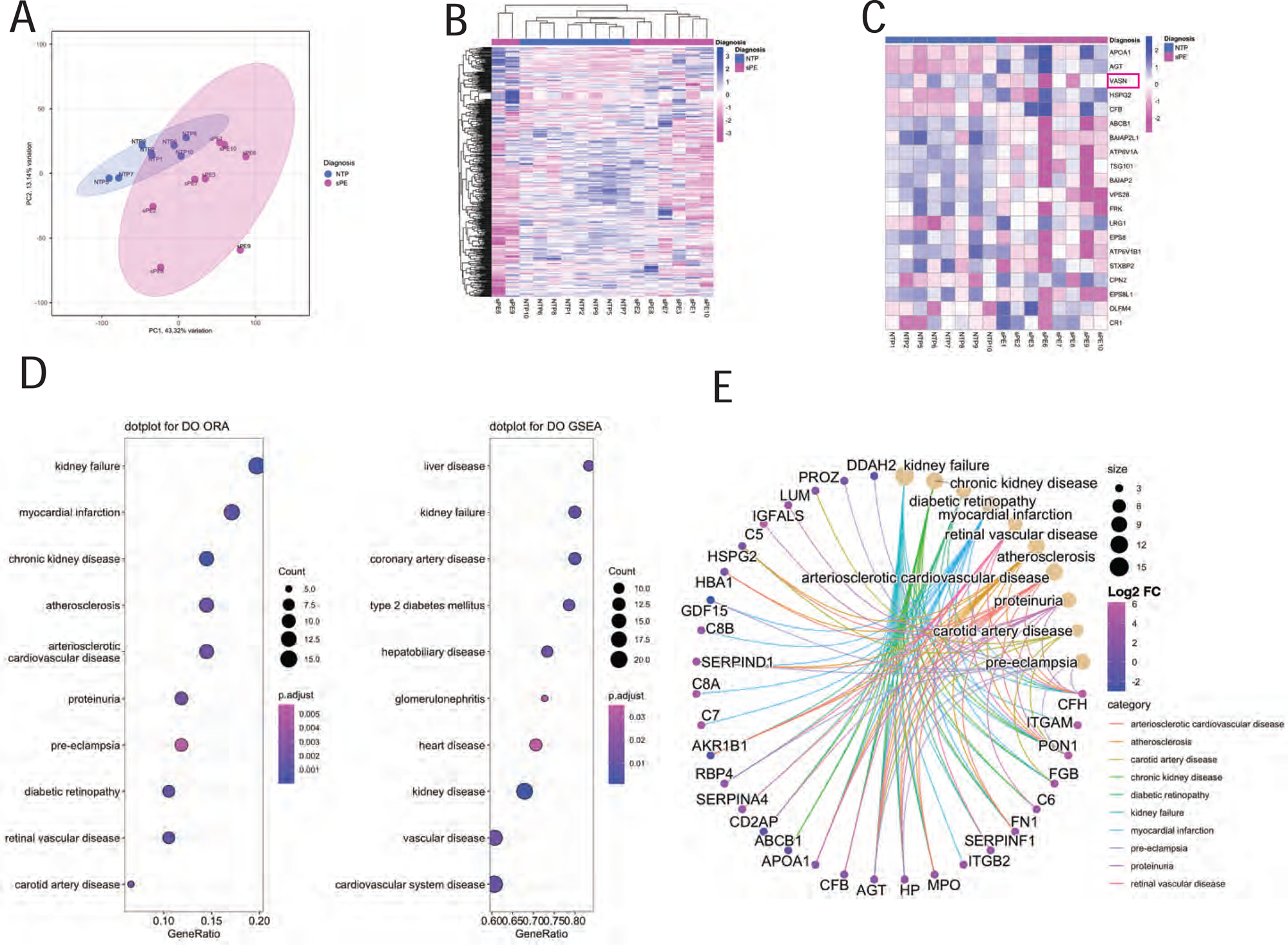
Unbiased proteomic analysis of urinary EV from NTP and sPE. Urine was collected form a cohort of women with NTP or sPE (n=10 NTP and n=10 sPE), EV were isolated and unbiased proteomic analysis was performed as described in methods. (**A)** shows results of principal component analysis (PCA) performed on normalized counts of all detected proteins in the samples. (**B)** illustrates a heatmap with hierarchical clustering performed on the same samples and detected proteins as were used for the PCA. Both PCA and hierarchical clustering indicate separation of the NTP and sPE groups and clustering of samples belonging to each group. **(C)** depicts a list of the top 20 significantly regulated proteins between the groups ranked in the order of absolute Z-scores. We highlighted VASN as the subject of our follow up study. (**D)** shows dot plots of disease ontology (DO) analysis using either over representation analysis (ORA; left) or gene set enrichment analysis (GSEA; right). (**E)** depicts gene concept network of the identified proteins as they relate to pathways shown in DO OR. Relative change of quantities of individual proteins is indicated by the color of the dot representing them; decrease (blue) or increase (magenta). The most significantly regulated DO pathways prominently feature disease processes related to vascular, kidney and hepatic diseases and include PE.

### Characterization EV, and VASN levels in plasma EV, total plasma, EV-depleted plasma and placenta from NTP and sPE Pregnant Women

**Figure 2** illustrates the isolation and characterization of Plex EVs obtained from both NTP and sPE patients. Transmission electron microscopy confirmed the presence of typical cup-shaped morphology (30–130 nm) associated with EVs in isolated Plex from pregnant women **(Figure 2A).** No significant differences were observed in the EV size distribution between the NTP and sPE groups **(Figure 2B).** Western blot analysis was performed to assess the purity and enrichment of EV in the isolated fractions. Samples included isolated plasma EV, whole plasma, HAEC lysates, and PBS as a negative control. Lanes were probed with specific antibodies targeting CD63 and TSG101 (enriched EV markers), GM130 (Golgi marker to exclude cellular contamination), and Apolipoprotein A1 (ApoA1, a plasma protein). The results **(Figure 2C)** demonstrate enrichment of CD63 and TSG101 in the EV fraction, while the EV-depleted plasma lacked these markers. Furthermore, the EV fraction lacked GM130, indicating minimal cellular contamination, and contained ApoA1, a marker present in plasma and which was also detected in urinary EV **(Figure 1C)**. Quantitative analysis confirmed the enrichment of VASN and TSG101 in the EV fraction and their depletion in the EV-depleted plasma **(Figure 2D).** Western blot analysis revealed decreased VASN protein levels in sPE plasma EVs compared to NTP EVs **(Figure 2E),** with TSG101 used for normalization as an EV marker. This pattern of decreased VASN protein was also observed in placental tissue from the sPE group compared to NTP **(Figure 2F),** suggesting potential depletion of VASN from plasma due to EV isolation. The presence of VASN within isolated plasma EVs was further confirmed by western blotting **(Figure 2 D, E).** RNA isolated from placenta samples demonstrated downregulation of VASN mRNA levels in the sPE group compared to the NTP group **(Figure 2H),** corroborating the findings at the protein level.

**Figure 2.**
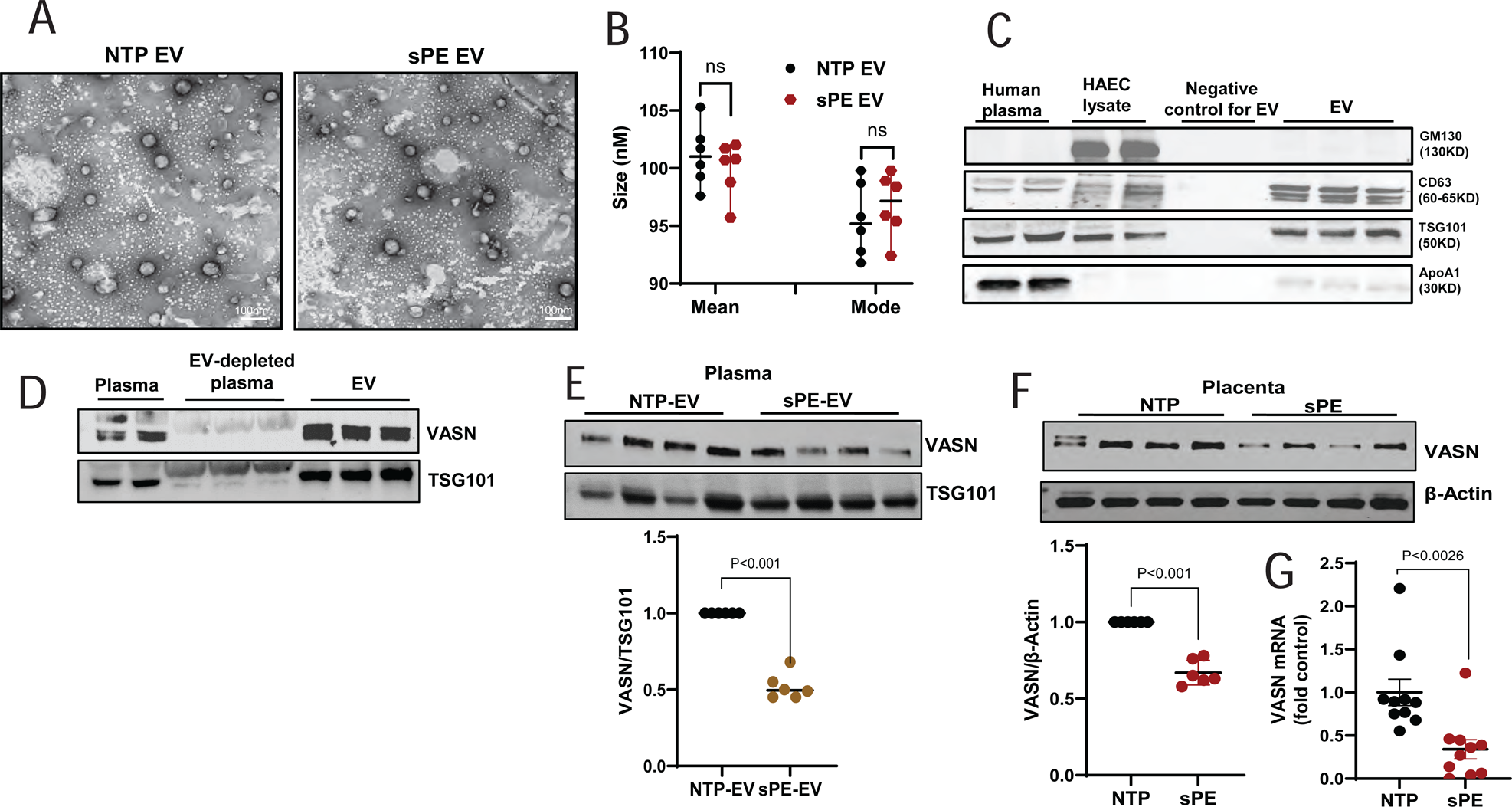
Characterization of EV, and VASN levels in plasma EV, total plasma, EV-depleted plasma and placenta from NTP and sPE Pregnant Women. EV were isolated from plasma of the NTP and sPE groups then (**A)** transmission electron microscopy (TEM) was performed to assess morphology. Scale bar: 100 nm. Both groups display typical EV morphology, characterized by cup-shaped structures within the size range consistent with EV. **(B)** Particle size for both EV groups were determined by Nanoparticle Tracking Analysis (NTA). No statistically significant difference (ns) was observed in size distribution between NTP and sPE EVs (t-test). (**C)**, We performed western blot analysis of isolated plasma EVs, along with whole plasma, HAEC lysates and PBS as a negative control for EV. Lanes represent samples probed with specific antibodies: CD63 and TSG101 (enriched EV markers), GM130 a Golgi marker to exclude cellular contamination and Apolipoprotein A1 (ApoA1) - a plasma protein. The EV fraction displays enrichment and the EV depleted plasma is void of the known EV markers (CD63 and TSG101), while lacks GM130 and contains Apo A1. **(D)** Quantitative analysis of VASN and TSG101 enrichment and depletion in EV and EV-depleted plasma, respectively. Quantitative analysis of VASN level changes by western blotting in sPE vs NTP plasma EV (**E)**, and placenta samples (**F)**, and by real time PCR in plasma samples **(G).** VASN protein is decreased in sPE compared to NTP EV, with TSG101 used as an EV marker for normalization. This pattern was also observed in placenta tissue **(F)**, suggesting potential VASN depletion from plasma by EV isolation. Western blotting confirmed the presence of VASN within isolated plasma EV **(G). (H)** RNA isolated from placenta samples demonstrated a downregulation of VASN mRNA levels in the sPE group compared to NTP. Data were presented as the mean ± SEM. Exact P values are shown as determined by Two-tailed, unpaired t-test.

### Levels of VASN in placenta tissue and plasma EV in normal murine pregnancy and in the sFLT-1 over-expression murine model of PE

To investigate the potential role of EV, and quantify VASN levels during pregnancy in a PE model, we employed a timed pregnant mouse model. Mice were injected intravenously with either adenovirus encoding human soluble fms-like tyrosine kinase-1 (AD-hsFLT-1) or a control adenovirus encoding enhanced green fluorescent protein (AD-eGFP) at embryonic day (E) 11.5. Blood and placenta samples were collected at subsequent time points: E15.5, E17.5, and E19.5. Successful model establishment was confirmed by detecting hsFLT-1 expression in the livers of AD-hsFLT-1 injected mice and eGFP expression in AD-eGFP injected mice **(Figure 3A, B).** Additionally, high levels of hsFLT-1 were verified in the plasma of AD-hsFLT-1 injected mice **(Figure 3C).** As expected, mice administered AD-hsFLT-1 exhibited significant fetal loss **(Figure 3D, E)** and displayed signs of liver damage, as indicated by elevated plasma alanine aminotransferase (ALT) levels **(Figure 3F).** In untreated (UT) pregnant mice, VASN levels progressively increased with advancing gestational age (GA) in both placental tissue and plasma EV **(Figure 3G, H).** However, in pregnant mice administered sFLT-1, VASN levels decreased with advancing GA in both placental tissues and EV **(Figure 3G, H).**

**Figure 3.**
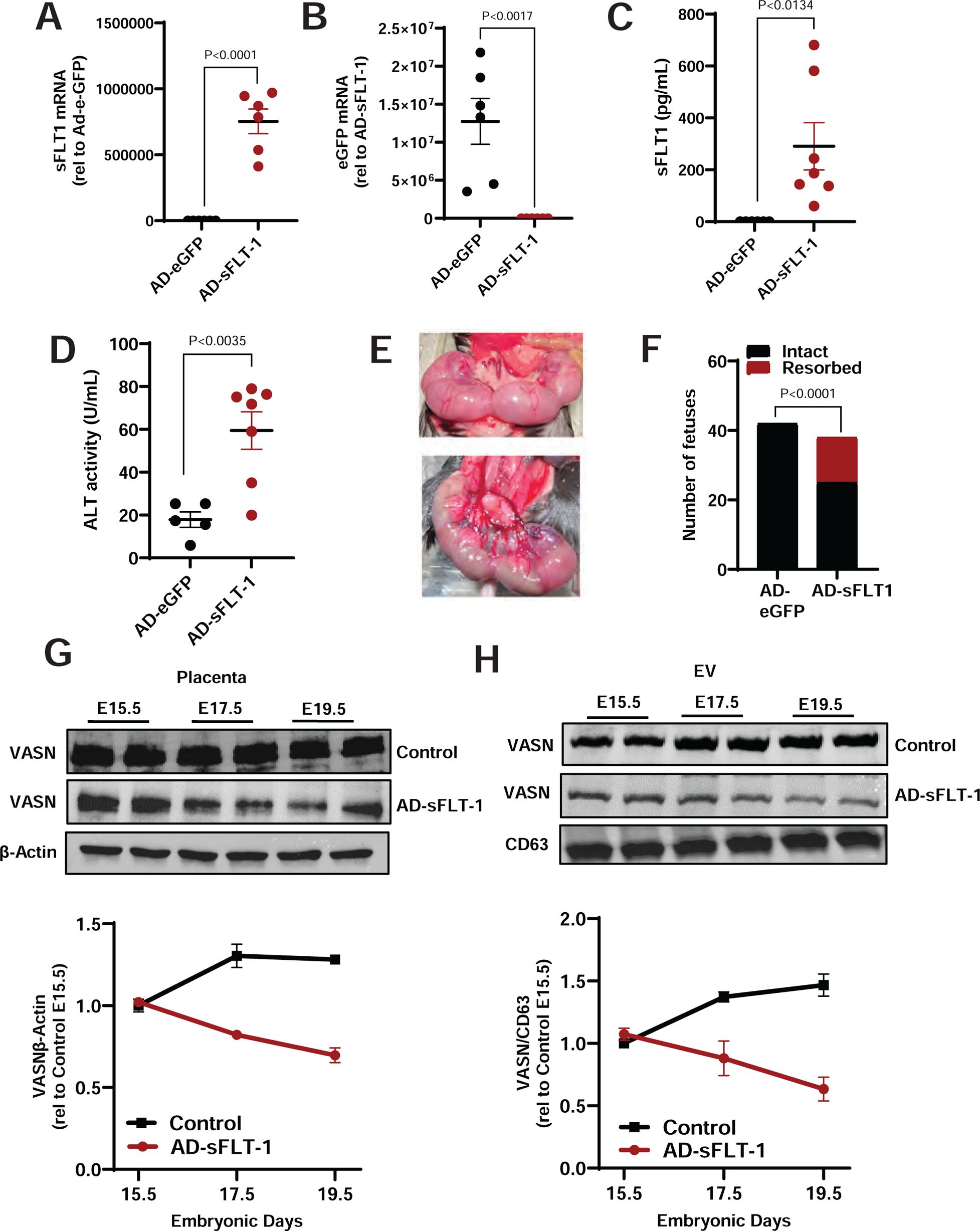
Levels of VASN in placenta tissue and plasma EV in normal murine pregnancy and in the sFLT-1 over-expression murine model of PE. Timed pregnant mice were injected intravenously with either AD-GFP (control) or AD-sFLT-1 (PE model) on E10.5 and on E15.5. sFLT1 mRNA (A) and eGFP mRNA (B) in liver, plasma and then we assessed on E18.5: **(A)** sFLT-1 mRNA (n=6), **(B)** GFP mRNA (n=6) in liver by real time PCR, **(C)** sFLT-1 protein (n=7) by ELISA, **(D)** alanine transaminase levels (n=7) by enzyme activity in plasma gross fetal morphology and survival **(E,F)** plasma. Administration of AD-sFLT-1 resulted in sFLT-1 mRNA expression in the liver and elevated sFLT-1 protein levels in the plasma, fetal loss and elevated plasma ALT level. We assessed on E15.5, 17.5 and 19.5: VASN levels in **(G)** placenta, **(H)** plasma EV (n=4 at each time point in each group). VASN levels exhibit gestational age (GA)-dependent increase in placenta and in plasma EV during normal murine pregnancy. Administration of AD-sFLT-1 resulted in decreased VASN in both placenta and in plasma EV. Data were presented as the mean ± SEM. Exact P values are shown as determined by Two-tailed, unpaired t-test.

### The roles of VASN in the regulation of vascular reactivity by sPE-EV

To test whether sPE-derived plasma EV can affect the regulation of vascular tone, and to test a potential role for VASN in this process, murine vascular rings (MVRs) were treated with EVs isolated from either NTP-EV or sPE-EV to assess their impact on endothelial function, then we assessed acetylcholine (ACh)-induced vasorelaxation in pre-constricted MVRs by wire myography. To determine if VASN is a potential mediator of these effects, we employed adenoviral vectors for VASN knockdown (AD-shVASN) and overexpression (AD-VASN). Incubation with sPE-EV for 24 hours significantly impaired ACh-mediated vasorelaxation **(Figure 4A, C).** Notably, pre-incubation with AD-shVASN alongside sPE-EV did not alter the vasodilatory response to ACh compared to treatment with sPE-EV alone **(Figure 4A).** Conversely, pre-incubation with sPE-EV followed by AD-VASN treatment restored the vasodilatory response to ACh, comparable to that observed with NTP-EV treatment **(Figure 4A).** Importantly, sodium nitroprusside (SNP)-induced relaxation remained unaffected by all treatments (NTP-EV, sPE-EV, AD-shVASN, or AD-VASN) **(Figure 4B, D).** Florescence microscopy images confirmed successful VASN overexpression in the murine aortic rings **(Figure 4E).** These findings suggest that VASN plays a critical role in endothelial dysfunction induced by sPE-EV. sPE-EV exposure impairs ACh-mediated vasorelaxation in MVR and this dysfunction can be reversed by increasing VASN expression.

**Figure 4.**
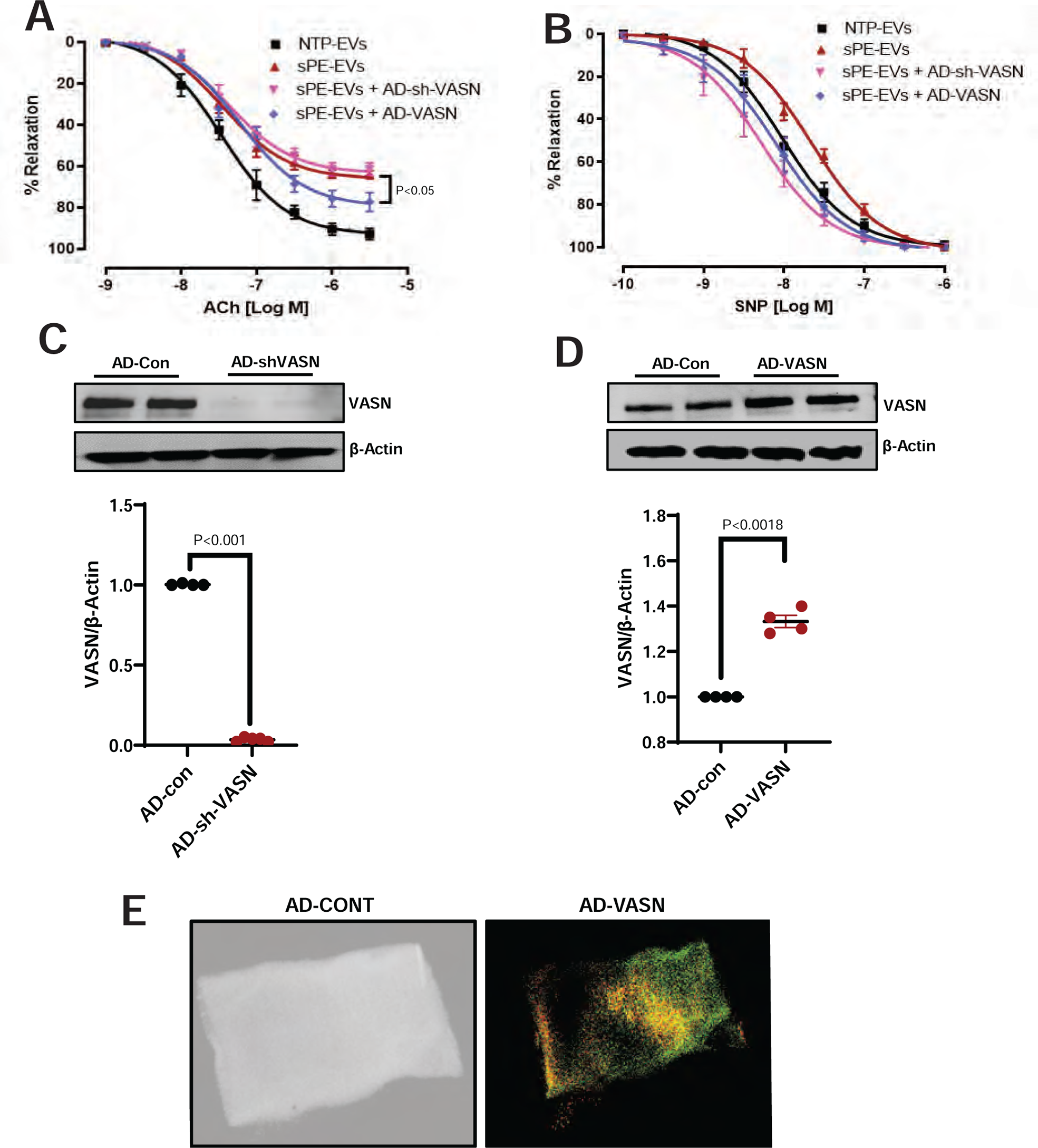
The roles of VASN in the regulation of vascular reactivity by sPE-EV. MVRs were prepared, exposed to AD-VASN or AD-shVASN overnight, and then were suspended in wire myograph chambers where they were maximally contracted by treatment with phenylephrine and then treated sequentially with acetylcholine (ACh) or with sodium nitroprusside (SNP) as indicated on the x axis. **(A)** depict the vasodilatory response of aortic rings to ACh, an endothelium-dependent vasodilator. **(B)** illustrate the response to SNP, an endothelium-independent vasodilator. Both ACh and SNP induced vasodilation (n=6 per condition). Western blot analysis confirmed VASN knockdown **(C)** using AD-sh-VASN and overexpression **(D)** using AD-VASN (graph represents n=4 per group). **(E)** Pre-treatment with AD-VASN significantly prevented the loss of ACh-induced vasorelaxation observed after sPE-EV treatment **(A).** SNP-induced vasodilation remained unaffected by all treatment groups **(B). (E)** Representative confocal images of VASN overexpression in the murine aortic rings. Data presented as mean ± SEM. Statistics by ANOVA with Tukey’s post-hoc test. Exact P values are shown as determined by Two-tailed, unpaired t-test.

### The effects of VASN knock down and over-expression on HAEC migration after treatment with sPE-EV

To investigate the role of VASN in sPE-EV-mediated inhibition of HAEC migration, we employed a scratch wound healing assay on HAEC monolayers with or without VASN manipulation using adenoviral vectors for overexpression (AD-VASN) and knockdown (AD-shVASN). Treatment with sPE-EV significantly impaired scratch wound closure compared to the control (NTP-EV), suggesting an inhibitory effect on HAEC migration **(Figure 5A, C).** The NTP-EV treatment itself did not affect migration. Knockdown of VASN (AD-shVASN) caused a slight but significant reduction in migration even in NTP-EV treated HAECs. Notably, AD-shVASN did not further decrease migration in cells treated with sPE-EV, suggesting VASN may not be the sole mediator of sPE-EV’s inhibitory effect **(Figure 5A, C).** Conversely, VASN overexpression (AD-VASN) completely reversed the inhibitory effect of sPE-EV on migration, restoring it to levels comparable to NTP-EV treated cells **(Figure 5A, C).** Importantly, treatment with a control adenovirus (AD-eGFP) had no effect on HAEC migration **(Figure 5B).**

**Figure 5.**
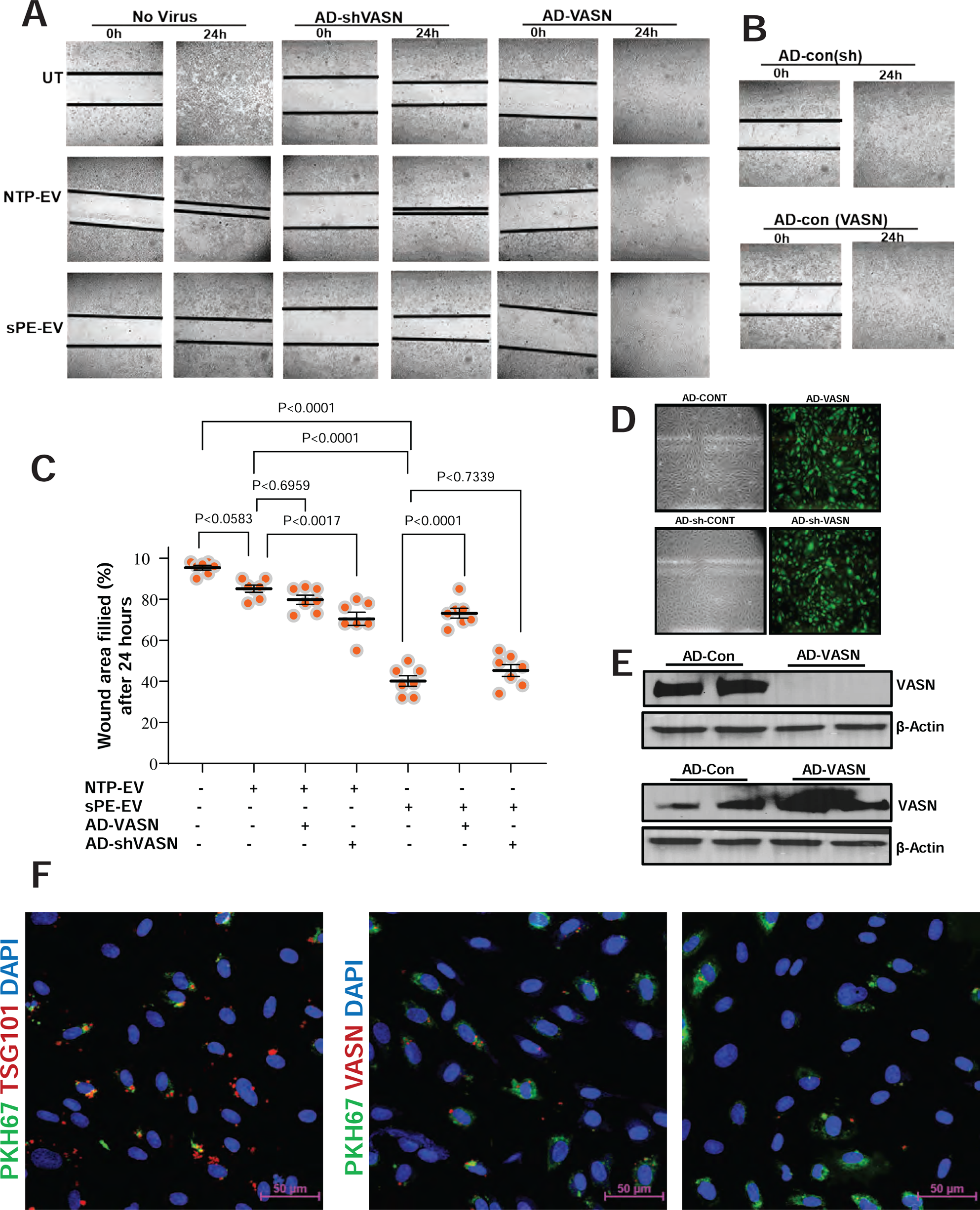
The effects of VASN knock down and over-expression on HAEC migration after treatment with sPE-EV. HAEC monolayers were untreated (UT) or were treated with AD-VASN and with AD-shVASN overnight, then were scratched and scratch wound closure was assessed 24 h later. **(A)** depicts representative images under each condition of virus/EV treatment. **(B)** effect of treatment with control (GFP) adenovirus. **(C)** summary graph with statistics (n=7 in each observation). **(D)** Assessment of transfection efficiency by visualizing eGFP fluorescence that is expressed in both AD-VASN and AD-shVASN from the bicistronic construct. **(E)** Western blot showing the effect of treatments on VASN expression. Treatment with sPE EV, but not with NTP EV inhibited HAEC migration into the wound. Over-expression of VASN significantly improved HAEC migration after treatment with sPE-EV. Knock down of VASN expression in HAEC with AD-shVASN inhibited HAEC migration in the presence of NTP EV and did not affect the migration of HAEC after sPE EV treatment. HAEC were treated with NTP-EV or with sPE-EV **(F)** that were labeled with an antibody to TSG101 (EV marker; red) or were labeled with an antibody against VASN (red) and with the lipophilic dye PKH67 (green). Nuclei were stained with DAPI (blue). Confocal images were collected and maximum intensity projections across four focal planes are shown. Robust uptake of EV was detected along with uptake of the EV marker and VASN. Data were presented as the mean ± SEM. Exact P values are shown, ANOVA with Tukey’s post hoc test.

Transfection efficiency was confirmed by visualizing eGFP fluorescence expressed in both AD-VASN and AD-shVASN due to the bicistronic construct design **(Figure 5D).** Western blot analysis documented the successful modulation of VASN expression by the adenoviral vectors **(Figure 5E).** To assess EV uptake by HAECs, cells were treated with either NTP-EV or sPE-EV labeled with antibodies against the EV marker TSG101 (red) or VASN (red), along with the lipophilic dye PKH67 (green) for membrane staining. Nuclei were visualized with DAPI (blue). Confocal microscopy revealed robust uptake of EV, co-localization with both the EV marker TSG101 and VASN **(Figure 5F).** These findings demonstrate that VASN plays a crucial role in EV-mediated inhibition of HAEC migration. While VASN knockdown partially mimics the inhibitory effect of sPE-EV, its overexpression completely abrogates it. Additionally, the co-localization of VASN with EVs suggests its potential role as a cargo molecule influencing migration.

### The effects of VASN over-expression or knock-down on gene expression changes in HAEC

To gain further insights into the mechanisms underlying VASN’s role in HAEC function, we performed RNA sequencing analysis on HAECs following VASN manipulation. Cells were untreated (control), treated with AD-VASN for overexpression, or treated with AD-shVASN for knockdown. After overnight incubation, RNA was isolated, and quality control (QC) was performed. RNA samples were then submitted for library preparation, sequencing, and subsequent analysis as described in the methods section. PCA revealed distinct clustering of the three treatment groups along both principal components 1 and 2 (PC1; x axis and PC2; y axis), indicating significant transcriptional alterations upon VASN modulation **(Figure 6A).** Separation along the PC1 axis appears to be related to the adenoviral transfection itself, while PC2 represents separation due to VASN over-expression or KO. Hierarchical clustering with a heatmap further confirmed this separation and demonstrated tight clustering within each treatment group **(Figure 6B).** A Venn diagram **(Figure 6C)** depicts the number of genes significantly regulated by VASN overexpression (magenta), VASN knockdown (blue), or by both treatments (light magenta). Volcano plots illustrate the log fold change and −log q values for differentially expressed genes, highlighting the most highly regulated genes **(Figure 6D, E)** by VASN over-expression **(Figure 6C)** and by VASN knockdown **(Figure 6E).**

**Figure 6.**
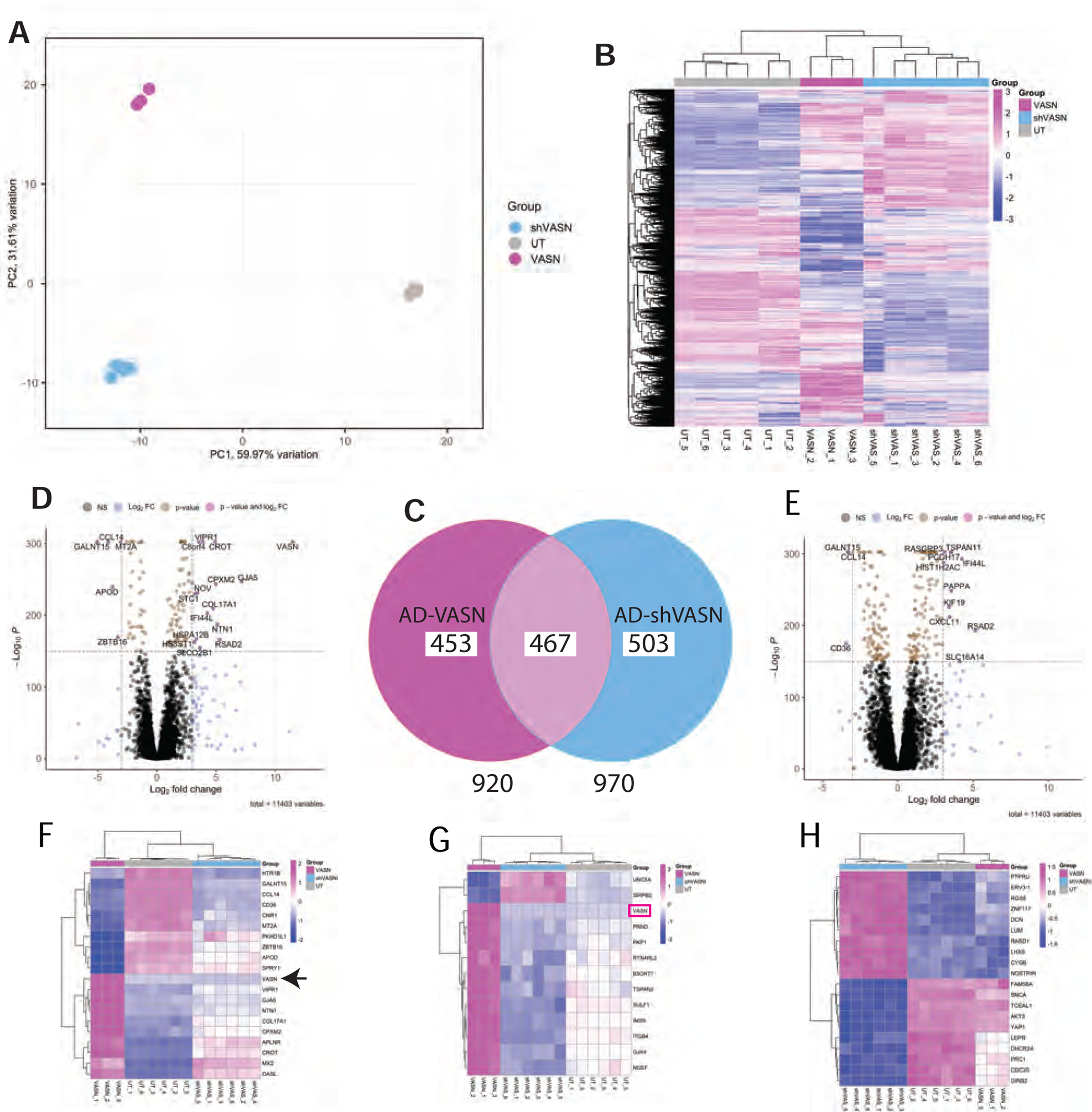
The effects of VASN over-expression or knock-down on gene expression changes in HAEC. Cells were untreated (UT) or were treated with AD-VASN and with AD-shVASN overnight, then RNA was isolated and QC was performed. RNA samples were submitted for library preparation and sequencing, then were analyzed as described in methods. PCA **(A)** and hierarchical clustering with heatmap **(B)** revealed that the three treatment groups exhibited high level of separation and the samples within groups exhibited tight clustering. Venn diagram **(C)** illustrates the number of genes that were significantly regulated only by VASN over-expression (magenta), only by VASN knock down (blue) or were regulated by both treatments (light magenta), along with volcano plots illustrating the log fold change and −log q values and most highly regulated genes **(D, E)**. **(F)** and **(H)** show the top 10 up and top 10 down regulated genes by VASN over-expression or knock-down, respectively. **(G)** depicts the 11 genes (including VASN) that were regulated in both VASN over-expression and knock-down, but in opposite directions.

Heatmaps of the top 10 upregulated and downregulated genes by VASN overexpression **(Figure 6 F)** and knockdown **(Figure 6 H)** are presented. Interestingly, 11 of the 467 genes that were regulated by both VASN over-expression and by VASN knockdown were found to be regulated in opposite directions **(Figure 6G).** These 11 genes included VASN itself as expected. The complete lists of differentially expressed genes, along with the relevant statistical data are presented in supplementary data. These findings indicate a complex role for VASN in regulating specific gene networks within HAECs.

### Pathway and network analysis of genes regulated by VASN over-expression and knock down in HAEC

Comparative over-representation analysis was performed to identify biological pathways and processes enriched by genes significantly regulated (fold change ≥ 2, q-value < 0.05) upon VASN overexpression or knockdown in HAECs. Data shown in figure are select pathways for each analysis that are related to known aspects of vascular function. The all-inclusive list of results for pathway analysis are shown in supplementary data. Results are presented in two formats: The color scale represents the significance of pathway activation, with warmer colors indicating higher significance. The size of each dot reflects the ratio of genes from the analysis participating in that specific pathway relative to the total number of genes in the pathway. Node sizes correspond to the ratio of participating genes in the pathway. Node and connecting line colors indicate the direction of regulation (up or down) for each gene in the four permutations (VASN overexpression/knockdown and upregulated/downregulated). **Figure 7 A and B** depict select DO pathways enriched by VASN-modulated genes. Notably enriched pathways include those related to (indicate specific pathways identified in the results). **Figure 7 C, D** show select KEGG pathways enriched by differentially expressed genes. This analysis revealed enrichment in pathways associated with (indicate specific pathways identified in the results). **Figure 7 E, F** depict results from the Reactome pathway analysis. This analysis identified significant enrichment in pathways involved in (indicate specific pathways identified in the results).

**Figure 7.**
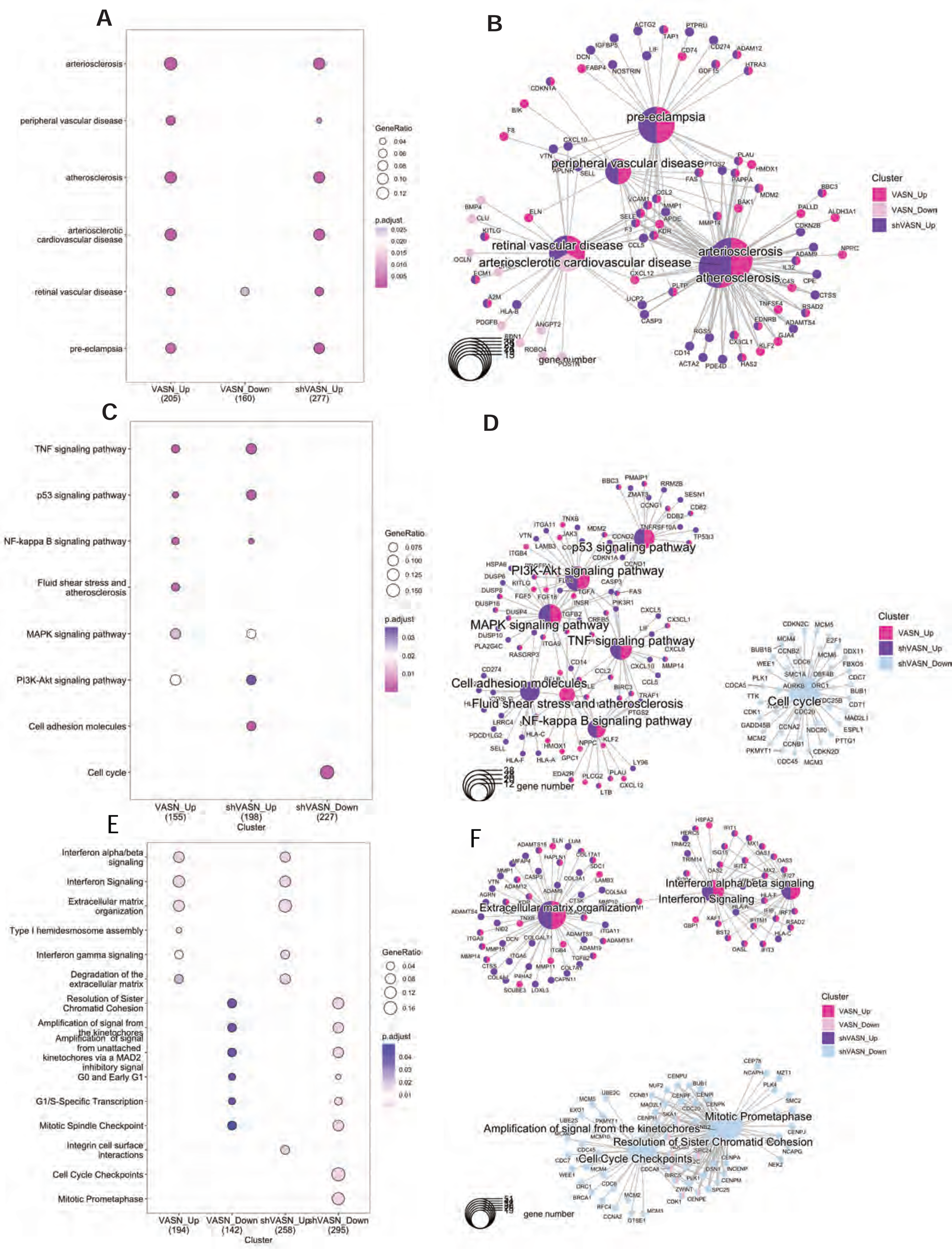
Pathway and network analysis of genes regulated by VASN over-expression and knock down in HAEC. Comparative gene over-representation analysis was performed using lists of significantly (2-fold change, q<0.05) up and down-regulated genes by VASN over-expression or knock-down in HAEC. Results are shown as dot blots **(A, C, E)**, where pathway activation significance is illustrated with the color scale shown and the ratio of participating genes relative to the total number of genes in the pathway is noted by the size of the dots, or as gene concept networks **(B, D, F)**, where the node sizes reflect on the ratio of participating genes and the color of the nodes and genes indicate whether they are regulated in any of the four permutations of VASN/shVASN/UP/DOWN. **(A, B)** illustrate select disease ontology (DO) pathways, panels **(C, D)** show select Kyoto encyclopedia of genes and genomes (KEGG) and panels **(E, F)** depict results of the reactome analysis.

### Characterization of placenta explant (Plex) derived extracellular vesicles (Plex-EV)

To establish an ex-vivo system to generate specifically placenta-derived EV, freshly collected placentas were processed for explant culture immediately after delivery. The supernatant and EV were isolated 24 hours later as described in the methods section. **Figure 8 A** shows the gross morphological appearance of explants under phase-contrast microscopy. **Figure 8 B** depicts the Western blot analysis that was performed to assess the purity and enrichment of EV in the isolated fractions. Samples included isolated plex EV, whole plasma, human placental tissue lysates, and PBS as a negative control. Lanes were probed with specific antibodies targeting CD63 and TSG101 (enriched EV markers), and Apolipoprotein A1 (ApoA1). The results (**Figure 2B**) demonstrate enrichment of CD63 and TSG101 in the Plex EV fraction. Furthermore, the EV fraction lacked ApoA1, indicating minimal or no cellular contamination. **Figure 8 C** presents TEM images of EV isolated from NTP and sPE placenta explants, closely resembling the morphology of plasma derived EV **(Figure 2A)**. **Figure 8 D** shows that the concentrations of Plex EV did not exhibit significant differences between the groups. **Figure 8 E** illustrates the size distribution of NTP and sPE placenta explant-derived EV. No significant differences were observed in the EV size distribution between the NTP and sPE groups.

**Figure 8.**
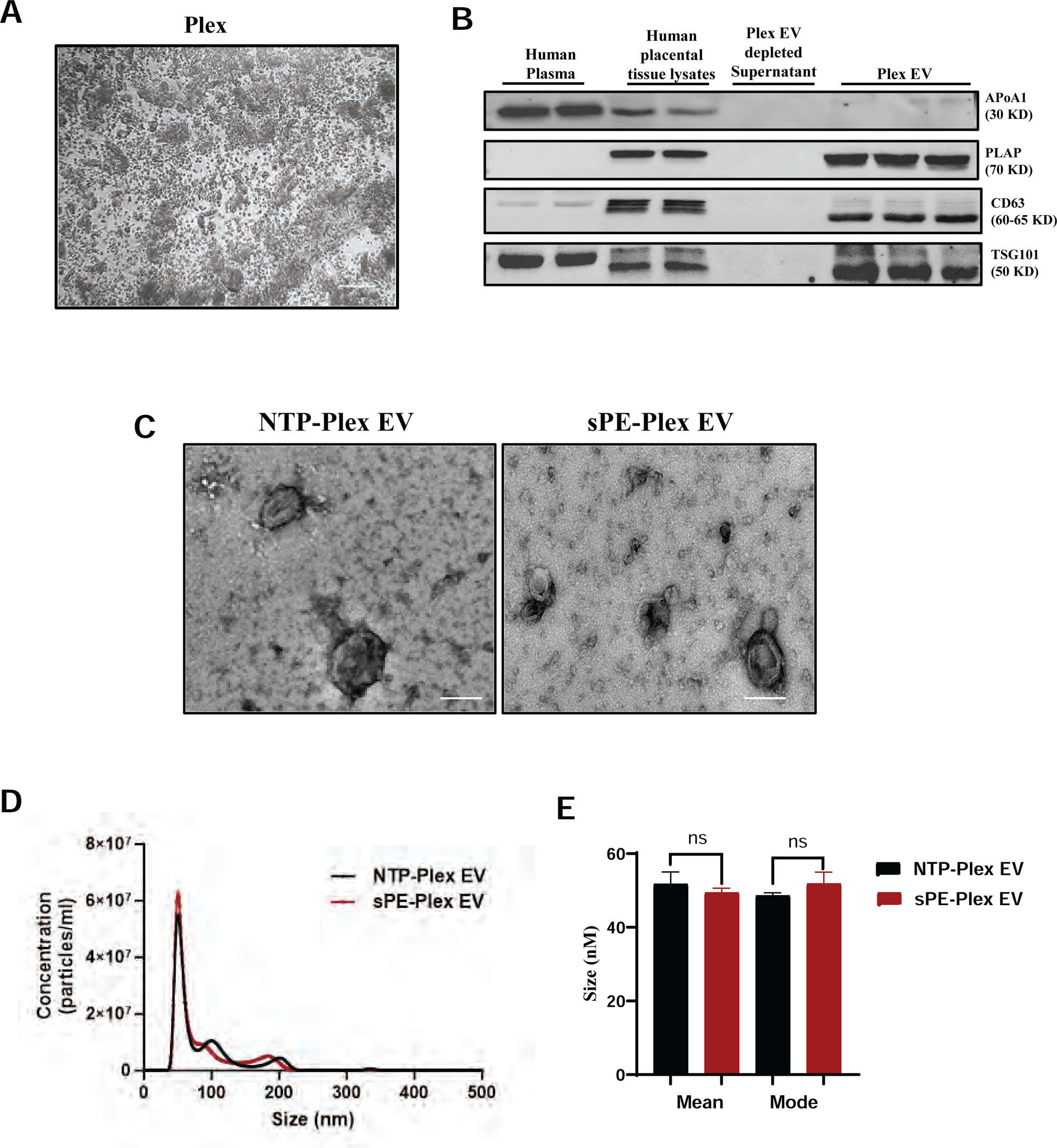
Characterization of placenta explant (Plex) derived EV (Plex-EV). Placentas were freshly collected after delivery and were processed to be cultured as placenta explants, then supernatant was collected 24h later and EV were collected as described in methods. **(A)** shows the gross morphological appearance of explants under phase contrast microscopy, Scale bar: 100 µm **(B)** illustrates distribution of ApoA1 plasma protein, placenta alkaline phosphatase (PLAP) placenta marker, CD63 and tumor suppressor gene 101 (TSG101) EV markers. **(C)** illustrates transmission electron micrographs of EV from NTP and sPE Plex. **(D)** shows particle size distribution of NTP and sPE Plex EV. Panel (E) is a summary graph of particle mean and mode size for NTP and sPE EV. Data were presented as the mean ± SEM. Exact P values are shown, Two-tailed, unpaired t-test.

### The effect of NTP-Plex-EV and sPE-Plex-EV on migration of HAEC in wound healing assays

To investigate the functional impact of sPE Plex EVs on HAEC, we employed a scratch wound healing assay on HAEC monolayers. HAEC monolayers were treated with either sPE Plex EV or NTP-Plex EV and control (without EV). Compared to the NTP-Plex EV treated cells, sPE-EV treatment resulted in a significant reduction in scratch wound closure, indicating impaired HAEC migration **(Figure 9A, B).** Notably, treatment with NTP-EV had no effect on migration, suggesting the observed effect is specific to sPE-EV and not a general consequence of EV exposure **(Figure 9A, C).** These findings demonstrate that sPE placenta explant-derived EVs possess an inhibitory effect on HAEC migration.

**Figure 9.**
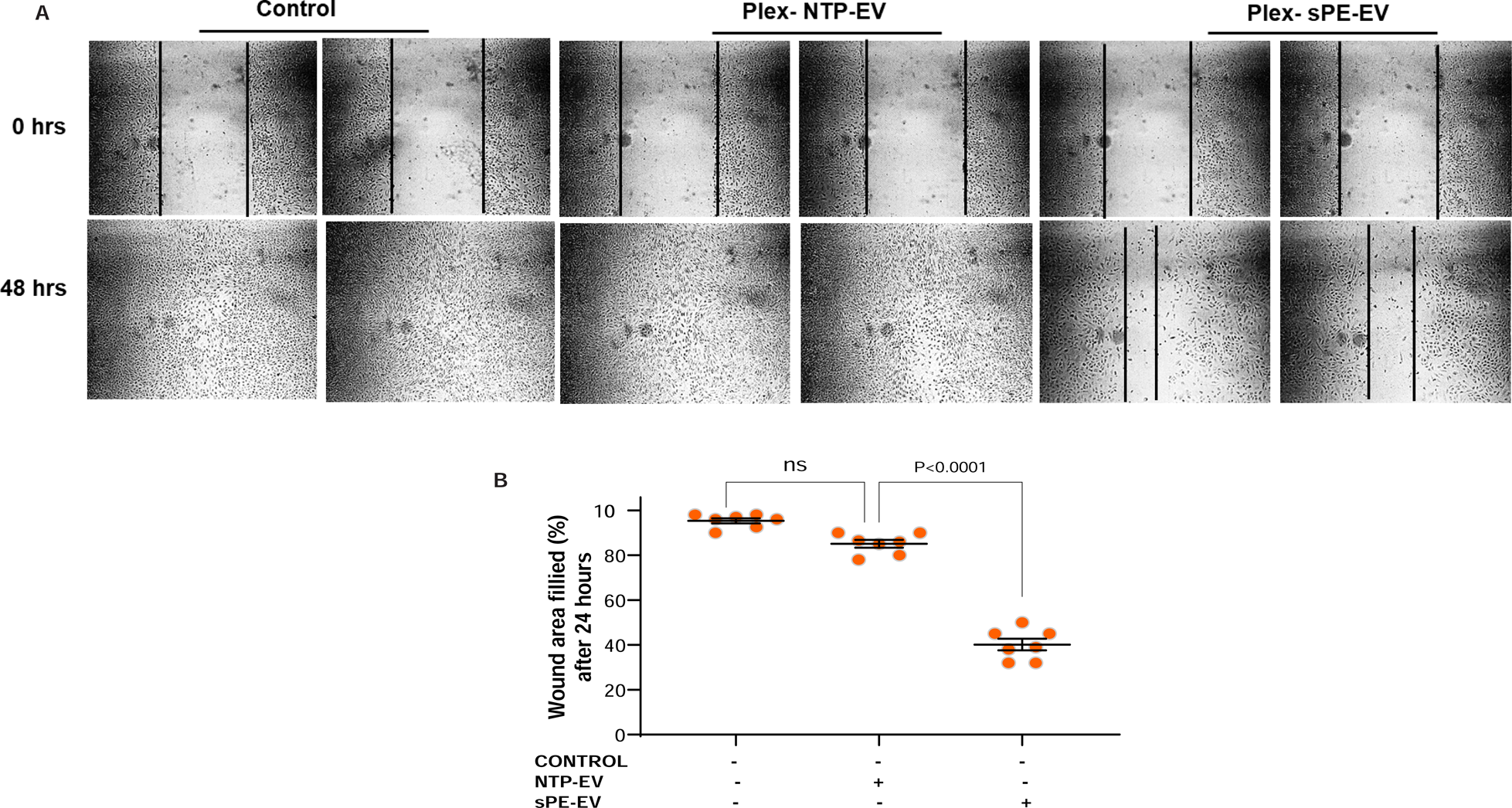
The effect of NTP-Plex-EV and sPE-Plex-EV on migration of HAEC in wound healing assays. Plex EV were prepared from explant cultures of NTP and sPE human placentas as described in methods, then scratch wound were created in confluent monolayers HAEC, followed by treatment with either NTP or sPE Plex-EV and analysis of wound closure 24h later **(A).** Wound closure was unaffected by treatment with NTP-Plex-EV, but was significantly inhibited by treatment with sPEPlex-EV**(B)**. Data were presented as the mean ± SEM. Exact P values are shown, ANOVA with Tukey’s post hoc test.

### The effect of EV isolated from murine placenta explant supernatants on HAEC migration in wound healing assays

In order to test whether we can generate EV from murine placenta explants that affect vascular function resembling the effects observed with plasma-derived EV, EV were isolated from explant cultures prepared from murine placentas obtained from mice injected with either control-AD-eGFP or AD-sFLT1. Confluent HAEC monolayers were subjected to scratch wounds and subsequently treated with the isolated Plex-EV. Wound closure was assessed after 24 hours**. Figure 10 A** shows the distribution of VASN and the EV marker TSG101 within the placental explants (Plex cultures), the EV-depleted supernatant, and the isolated EV. VASN is highly enriched in the isolated Plex-EVs compared to the unprocessed Plex cultures or the EV-depleted supernatant. VASN levels are significantly lower in the Plex-EV derived from sFLT1-injected mice (AD-sFLT1) compared to those from control mice (AD-eGFP) **(Figure 10B). >Figure 10C** presents representative images from the wound healing assay, illustrating the extent of scratch closure in HAEC monolayers treated with different conditions. Treatment with sFLT1-derived Plex-EVs (AD-sFLT1) significantly inhibits the migration of HAECs in the wound healing assay **(Figure 10 C, D).**

**Figure 10.**
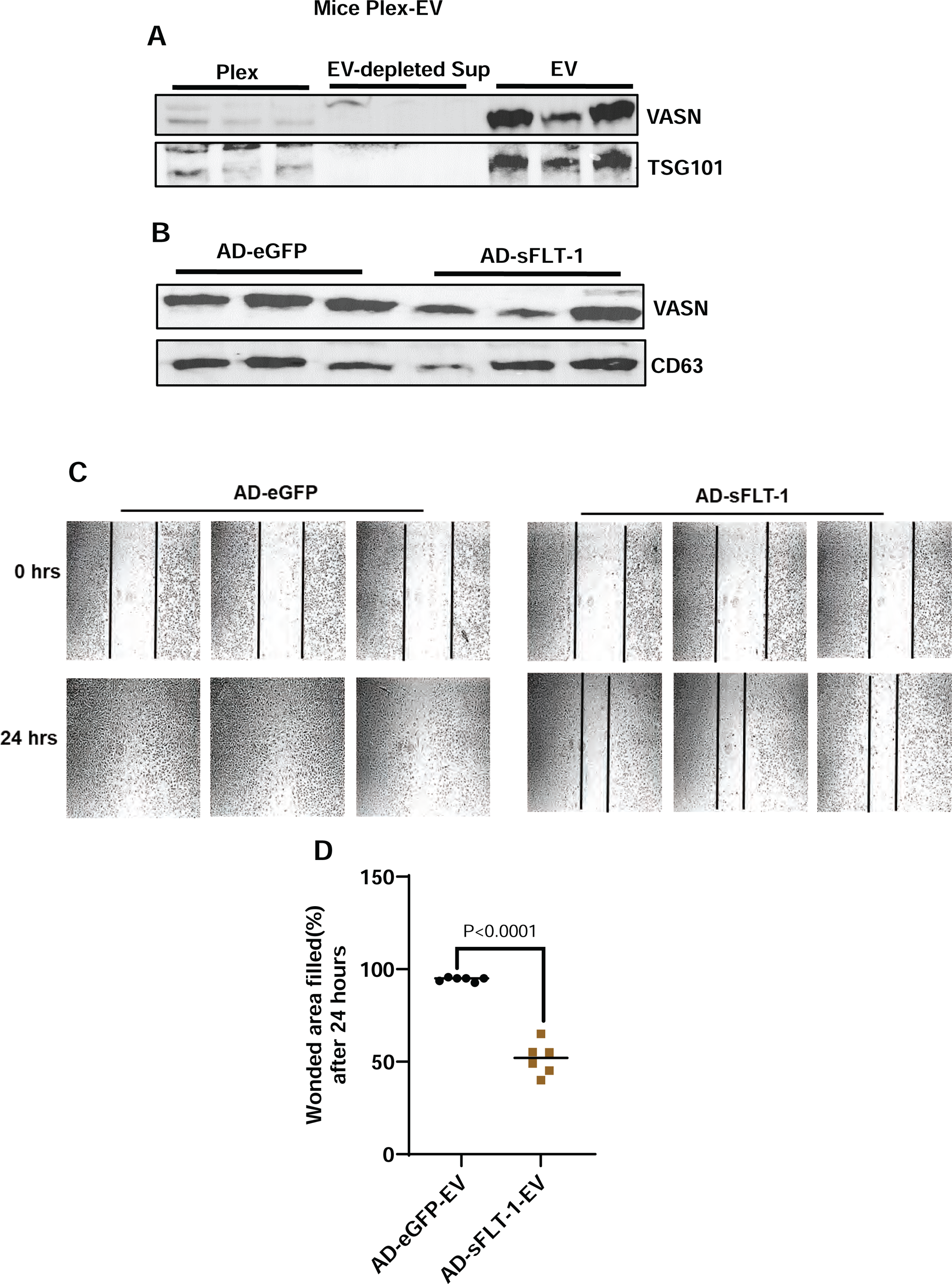
The effect of EV isolated from murine placenta explant supernatants on HAEC migration in wound healing assays. EV were isolated from explant cultures prepared from murine placentas from mice injected with adenovirus encoding GFP (control) or adenovirus encoding sFLT1 (murine model of PE). Scratch wound was made in confluent monolayers of HAEC, then they were treated with Plex-EV and scratch wound closure was assayed 24h later. Panel **(A)** documents the distribution of VASN and the EV marker TSG101 in Plex cultures, in EV-depleted supernatant and in EV. Panel **(B)** illustrates VASN levels in Plex-EV, as compared to the EV marker CD63 prepared from AD-eGFP and AD-sFLT-1 injected mice. Panel **(C)** shows representative images of the wound healing assay. Panel **(D)** is a graph of summary findings in wound healing assays. VASN is highly enriched in Plex EV as compared to Plex culture or EV-depleted Plex supernatant. VASN levels are decreased in AD-sFLT-1 Plex EV as compared to AD-eGFP Plex-EV. AD-sFLT1 Plex-EV significantly inhibits migration of HAEC. Data were presented as the mean ± SEM. Exact P values are shown, Two-tailed, unpaired t-test.

## DISCUSSION

Previously, we observed a significant increase in plasma EV content in sPE and demonstrated that sPE EV treatment induced endothelial dysfunction *in vitro*^21^. To identify candidate molecules that may mediate this effect, we conducted unbiased proteomics using EV isolated from urine samples of NTP and sPE, avoiding non-EV contaminants. Urinary EV, commonly used in biomarker discovery for kidney and prostate pathology, also reflect on disease states in various organs outside the urogenital system, including complications of pregnancy^47–51^. EV and exosomes during normal pregnancy and in PE have been extensively studied with over two hundred articles listed in PubMed; recently reviewed by Paul et al^52^. The majority of publications identifying specific vesicle cargo with an impact on pregnancy focused on miRNA content, with only a handful concerned with the proteome^53–58^. Longitudinal studies of gestational age-dependent changes of EV cargo has been restricted to miRNA^59–61^. Our study is unique, in that we identified candidate protein cargo with potential mechanistic significance in PE. Moreover, we verified in both human samples and in a murine model that VASN, the candidate that we identified, is decreased in sPE in humans and decreased in sFLT-1 over-expressing mice (a recognized murine model of PE^62^). Furthermore, we performed mechanistic testing by over-expression and knock down of VASN in the target cells or tissues, revealing a vascular endothelial phenotype that is dependent on levels of VASN expression.

We found that in pregnant mice VASN levels increase during the course of gestation in the placenta as well as in plasma EV. In mice injected with AD-sFLT-1, the trend reverses, and VASN decreases during gestation. In these respects, VASN exhibits similar tendencies during pregnancy as PlGF, which also increases with gestion in humans until the 32^nd^ week of pregnancy and decreases in PE. PlGF is a VEGFR1 agonist, which also activates VEGFR1/VEGFR2 heterodimer, resulting in signaling that is pro-angiogenic, and also acts on immune cells ^63,64^. Similar to PlGF, VASN plays regulatory roles in vascular function which are consistent with an overall pro-angiogenic effect: 1) VASN expression in endothelial cells, or exposure of endothelial cells to conditioned media collected from VASN over-expressing cells promotes angiogenesis in vitro^65,66^, 2) VASN knock out mice exhibit impaired NO-dependent vasorelaxation^67^, 3) VASN antagonizes vascular wall ageing^39^. Some of these effects have been attributes to antagonism of TGF-β signaling and/or direct activation of VEGFR2 by VASN^37,39,66,68^. All of these pro-angiogenic effects are consistent with the signaling needs of the developing placenta and vascular adaptation in normal pregnancy.

Indeed, our findings agree with the notion that VASN plays a pro-angiogenic effect. Treatment with sPE plasma EV with reduced VASN content caused changes in human aortic endothelial cells (HAEC) and murine vascular rings consistent with PE-induced anti-angiogenic effects and endothelial dysfunction. This effect is likely due to the combined contribution of a number of EV cargo molecules, including miRNA-s and proteins^32,34,69,70^. Notably, overexpression of VASN alone was able to reverse the effects of sPE EV on both isolated vessels and on endothelial cells in culture, suggesting that VASN plays a mechanistic role to block a critical common step in the anti-angiogenic effect by sPE EV content.

RNA-seq analysis performed on HAEC after over-expression and knock down of VASN provided additional insight into VASN-s role in mediating endothelial function. While a large number of genes were regulated in response to both VASN over-expression and knock down, approximately half of all regulated genes were similarly up or downregulated in both treatment groups. However, we identified 11 transcripts that were significantly regulated in both treatments, but in opposite directions. VASN was in this set, regulated +2865 fold (q<10^−100^) in AD-VASN-treated HAEC and regulated −2.18 fold (q=5.39^−31^) in AD-shVASN transfected HAEC, confirming the expected over-expression and knock down. There were only two genes that were significantly upregulated in AD-shVASN treated cells and were significantly downregulated in AD-VASN treated cells (UNC5A and SIRPB2). UNC5A is a receptor for netrin, which is primarily known for its role in axon guidance, but a pro-angiogenic role has been shown in endothelial cells^71,72^. SIRPB2 (signal regulatory protein beta) is a poorly understood gene, but has been noted to be regulated by shear stress in endothelial cells^73^. Among the genes that were significantly downregulated in AD-shVASN treated cells and were significantly upregulated in AD-VASN treated cells included PRND, PKP1, TSPAN2 and SULF1. PRND is a prion protein with known angiogenic roles^74^, PKP1 is a desmosome component that plays a role in cell migration^75^, TSPAN2 is a tetraspanin receptor that is highly regulated by TGF-β and is known to reguate endothelial cell migration and proliferation^76^ and SULF1 is a heparan sulfate sulfatase that regulates the activities of heparan sulfate-binding VEGF isoforms^77^. Disease ontology analysis identified several vascular disease pathways that were affected by manipulation of VASN expression, including arteriosclerosis, retinal and peripheral vascular disease, and even PE was among the most affected disease pathways. Among the KEGG pathways that were affected by VASN KO and over-expression notable are the MAPK and PI3K/Akt signaling pathways that are known regulators of endothelial cell migration, the NFkB pathway that is critical in both inflammation and cell survival and the TNF pathway. Notably, the gene sets that were inhibited by VASN knock down appear to be all related to cellular proliferation and survival. These data provide string indication that VASN is a critical regulator of endothelial cell functions.

Our study has limitations. The PE cohort has a higher BMI compared to the NTP group, potentially confounding the observed changes in VASN levels. Future studies should include BMI-matched cohorts to isolate the independent effect of obesity on VASN. Additionally, the PE cohort was restricted to late-onset sPE, and did not cover the full spectrum of PE phenotypes. We focused on late onset cases, due to higher prevalence and easier recruitment. We decided to contrast PE with severe features to NTP to maximize effect size, expecting that vascular mediators with functional significance altered more in sPE than in PE without severe features. Future work will assess VASN levels in both early and late-onset PE with and without severe features, along with gestational age-matched NTP controls. The limitations of our strategy of human subject selection are partially mitigated by our findings in the murine model. We utilized mice with identical genetic backgrounds and weights, allowing for controlled comparisons. The model also demonstrated both gestational age-dependent changes in VASN levels and a decrease in VASN levels mirroring our observations in human sPE compared to NTP.

Taken together, our data suggest that reduced VASN in EV are potential biomarkers and functional drivers of endothelial dysfunction in PE. VASN has been implicated in the regulation of vascular function, but to our knowledge this is the first publication to; 1) provide a detailed account of GA-dependent changes of VASN in murine pregnancy; 2) demonstrate a PE-dependent change of VASN in human samples; 3) document altered VASN levels in a murine model of PE; 4) demonstrate that EV-encapsulated VASN can be produced from both human and murine Plex; 5) show that the VASN content of EV isolated form Plex reflects on the physiological state of Plex donors, i.e., whether the human subject had sPE, or the mouse was injected with sFLT-1 to induce a PE-like physiological state and 6) report on functional consequences of EV VASN on endothelial cells. Our findings suggest that quantitative changes in VASN may serve both as a biomarker and that VASN is a functional molecule that prevents/antagonizes endothelial dysfunction. Indeed, VASN may be a critical placental mediator of the cardiovascular adaptive response to pregnancy and it’s dysregulation may play an important role in the sPE maladaption. Further research on VASN may lead to diagnostic and therapeutic advancements improving outcomes in PE.

## Acknowledgments

The authors are grateful to High-Resolution Imaging Facility (HRIF, UAB) for providing access to the Confocal Microscopy. This work was supported by grants of Reinvent Program in the Department of Anesthesiology at UAB (MFP, SM), by R21ES031559 (TJ) and K23HL159331 (RGS).

## Contributions

SM provided design the study, performed, analyze the data, and write the manuscript; DA supported PE mouse experiment’s and sFIt-1 adenovirus; HH provided the clinical sample support and patient enrollment; MP helped the all-clinical aspect and patient enrollment, IRB; LS help the most of the experiments and data analysis; AS helped the patients samples collections; RS provided the clinical PE guidance and manuscript review and support; MT helped the patient enrollment, IRB; ZM helped the PE mice experiments; JM helped the proteomics experiment’s and analysis the data; AT provide the clinical support and samples collections and pregnancy patients guidance; TJ helped design the study, performed, analyze the data, and write the manuscript; DB provided design the study, analyze the data, and write the manuscript.

## Conflict-of-interest disclosure

The authors declare no competing financial interests.

## Nonstandard Abbreviations and Acronyms

PE: Preeclampsia
sPE: severe Preeclamptic pregnancies
EV: Extracellular Vesicles
NTP: Normotensive Pregnant Women
NTP-Plex EV: Normotensive Placental EV
sPE Plex EV: severe Preeclamptic Placental EV
HAEC: Human Aortic Endothelial Cells
Plex: Placental explant culture
NTA: Nanoparticle Tracking Analysis
TEM: Transmission electron Microscopy
MVRs: Murine vascular rings

---

**Supplemental Table 1:** Clinical Patient Characteristics.

| **Demographics** | **NTP (n=15)** | **sPE (n=15)** |
| --- | --- | --- |
| Maternal age | 30.3 ±1.2 | 28.6 ±1.6<sup>ns</sup> |
| Gestational age at delivery (week) | 38.12±0.5 | 36.65±0.8<sup>ns</sup> |
| BMI (kg/m2) | 30.96±1.5 | 45.2±2.1<sup>P<0.0001</sup> |
| Systolic blood pressure (mm/Hg) | 118.5 ±1.56 | 174.9 ±1.75<sup>P<0.0001</sup> |
| Diastolic Blood Pressure ((mm/Hg) | 71.25 ±1.6 | 100.13±3.6<sup>P<0.0001</sup> |
| Protein: creatinine ratio | ----- | 2.89±0.68<sup>P<0.0001</sup> |

Values are presented as mean ± SEM. All pregnancies were singleton without intrauterine infection or any other medical condition. P< 0.0001 vs NTP. Abbreviations: BMI, body mass index; BP, blood pressure; n.s, not significant; SEM, standard error of the mean; sPE, preeclampsia with severe features.

## Notes

### Competing Interest Statement

The authors have declared no competing interest.

